# Identification of NAD-dependent xylitol dehydrogenase from *Gluconobacter oxydans* WSH-003

**DOI:** 10.1101/634238

**Authors:** Li Liu, Weizhu Zeng, Guocheng Du, Jian Chen, Jingwen Zhou

**Author notes:** Corresponding author: Jingwen Zhou Mailing address: School of Biotechnology, Jiangnan University, 1800 Lihu Road, Wuxi, Jiangsu 214122, China.

## Abstract

*Gluconobacter oxydans* plays important role in conversion of D-sorbitol to L-sorbose, which is an essential intermediate for industrial-scale production of vitamin C. In the fermentation process, some D-sorbitol could be converted to D-fructose and other byproducts by uncertain dehydrogenases. Genome sequencing has revealed the presence of diverse genes encoding dehydrogenases in *G. oxydans*. However, the characteristics of most of these dehydrogenases remain unclear. Therefore, analyses of these unknown dehydrogenases could be useful for identifying those related to the production of D-fructose and other byproducts. Accordingly, dehydrogenases in *G. oxydans* WSH-003, an industrial strain used for vitamin C production, were examined. An NAD-dependent dehydrogenase, which was annotated as xylitol dehydrogenase 2, was identified, codon-optimized, and expressed in *Escherichia coli* BL21 (DE3) cells. The enzyme exhibited high preference for NAD^+^ as the cofactor, while no activity with NADP^+^, FAD, or PQQ was noted. Although this enzyme presented high similarity with NAD-dependent xylitol dehydrogenase, it showed high activity to catalyze D-sorbitol to D-fructose. Unlike the optimum temperature and pH for most of the known NAD-dependent xylitol dehydrogenases (30°C–40°C and about 6–8, respectively), those for the identified enzyme were 57°C and 12, respectively. The *K*_*m*_ and *V*_*max*_ of the identified dehydrogenase towards L-sorbitol were 4.92 μM and 196.08 μM/min, respectively. Thus, xylitol dehydrogenase 2 can be useful for cofactor NADH regeneration under alkaline conditions or its knockout can improve the conversion ratio of D-sorbitol to L-sorbose.

**Importance:** Production of L-sorbose from D-sorbitol by *Gluconobacter oxydans* is the first step for industrial scale production of L-ascorbic acid. *G. oxydans* contains a lot of different dehydrogenases, among which only several are responsible for the conversion of D-sorbitol to L-sorbose, while others may responsible for the accumulation of byproducts, thus decreased the yield of L-sorbose on D-sorbitol. Therefore, a new xylitol dehydrogenase has been identified from 44 dehydrogenases of *G*. *oxydans*. Optimum temperature and pH of the xylitol dehydrogenase are different to most of the known ones. Knock-out of the dehydrogenase may improve the conversion ratio of D-sorbitol to L-sorbose. Besides, the enzyme exhibits high preference for NAD^+^ and have potential to be used for cofactor regeneration.

## Introduction

The genus *Gluconobacter* is a part of the group of acetic acid bacteria, which are characterized by their ability to incompletely oxidize a wide range of carbohydrates and alcohols (1). *Gluconobacter* strains have been successfully used for the industrial production of food-related products, pharmaceuticals, and cosmetics, such as vitamin C (2), miglitol (3), dihydroxyacetone (DHA) (4), and ketogluconates (5). In particular, *Gluconobacter oxydans* has applications in the production of food additives and sweeteners owing to its ability to synthesize flavoring ingredients from aromatic alcohols, aliphatic alcohols, and 5-ketofructose (6, 7). Besides, *G. oxydans* enzymes, cell membranes, and whole cells are also used as sensor systems for the detection of polyols, sugars, and alcohols (8–10). In recently years, some *G. oxydans* strains have been employed for the production of enantiomeric pharmaceuticals and platform compounds; for example, *G. oxydans* DSM2343 has been employed for the reduction of various ketones used in pharmaceutical, agrochemical, and natural products (11), *Gluconobacter* sp. JX-05 has been utilized for D-xylulose and xylitol production (12), and *G. oxydans* DSM 2003 has been used for 3-hydroxypropionic acid production (13). As all of these products are related to the dehydrogenases of *G. oxydans*, identification of these enzymes in *G. oxydans* can expand the application of this bacterium.

*Gluconobacter* strains possess numerous dehydrogenases, some of which have been identified, such as alcohol dehydrogenase that could convert ethanol to acetaldehyde (14, 15), NADP-dependent acetaldehyde dehydrogenase that could convert acetaldehyde to acetate (16), PQQ-dependent glucose dehydrogenase that could convert D-glucose to D-gluconate (14), gluconate dehydrogenase that could convert D-gluconate to 2- or 5-ketogluconate (17), 2-ketogluconate dehydrogenase that could convert 2-ketogluconate to 2,5-diketogluconate (14), D-sorbitol dehydrogenase that could convert D-sorbitol to L-sorbose or fructose (14, 18–21), sorbose/sorbosone dehydrogenase that could convert L-sorbose to L-sorbosone or 2-KLG (22, 23), mannitol dehydrogenase that could convert mannitol to fructose (24, 25), quinate dehydrogenase that could convert quinic acid to shikimic acid (26–28), glycerol dehydrogenase that could convert glycerol to DHA (28), etc. In 2005, the complete genome of *G. oxydans* 621H was sequenced (29), which revealed 75 open reading frames (ORFs) that encode putative dehydrogenases/oxidoreductases of unknown functions. Identification of the functions of these unknown dehydrogenases/oxidoreductases is important to expand the application of *G. oxydans*. For instance, carbonyl reductase (GoKR) from *G. oxydans* DSM2343 has been employed for the reduction of various ketones (11), and membrane-bound alcohol dehydrogenase (mADH) and membrane-bound aldehyde dehydrogenase from *G. oxydans* DSM 2003 have been employed for 3-hydroxypropionic acid production (13).

In *G. oxydans*, the central metabolic pathway, such as citrate cycle and Embden-Meyerhof-Parnas pathway (EMP), is incomplete because of the absence of some genes encoding succinate dehydrogenase, phosphofructokinase, phosphotransacetylase, acetate kinase, succinyl-CoA synthetase, succinate dehydrogenase, isocitratelyase, and malate synthase (30, 31), which may be the reason for the low biomass of *G. oxydans* when cultured in rich medium. In a previous study, *sdhCDABE* genes encoding succinate dehydrogenase and flavinylation factor SdhE, *ndh* gene encoding a type II NADH dehydrogenase, and *sucCD* from *Gluconacetobacter diazotrophicus* encoding succinyl-CoA synthetase were expressed in *G. oxydans* to increase its biomass yield (31). However, *G. oxydans* biomass only increased by 60%, suggesting the presence of some unknown bottleneck. Except for the TCA cycle, all the genes were identified to encode enzymes involved in oxidative pentose phosphate and Entner-Doudoroff (ED) pathways (29). The pentose phosphate pathway is believed to be the most important route for phosphorylative breakdown of sugars and polyols to CO_2_ and provide carbon skeleton. It has been speculated that *G. oxydans* has the capability to uptake and channelize several polyols, sugars, and sugar derivatives into the oxidative pentose phosphate pathway; however, the gene involved in this process is still unknown. Hence, in the fermentation of sorbitol to sorbose for vitamin C production, some sorbitol must get converted to fructose or other byproduct to enter the pentose phosphate pathway for cell growth. Therefore, it is crucial to balance and control the conversion of sorbitol to fructose for cell growth and sorbose production.

Gene disruption and complementation experiments are often used to verify one gene function. Some *G. oxydans* genes have been identify by using this method, such as PQQ-dependent D-sorbitol dehydrogenase responsible for the oxidation of 1-(2-hydroxyethyl) amino-1-deoxy-D-sorbitol to 6-(2-hydroxyethyl) amino-6-deoxy-L-sorbose, which is the precursor of an antidiabetic drug miglitol (3), pyruvate decarboxylase that catalyzes the conversion of pyruvate to acetaldehyde by decarboxylation (32), mADH, membrane-bound inositol dehydrogenase, membrane-bound PQQ-dependent glucose dehydrogenase, etc. (33, 34). However, some *G. oxydans* genes encoding dehydrogenases are necessary for cell growth, and their knockout resulted in absence of growth (unpublished data). Besides, *G. oxydans* comprises numerous dehydrogenases, some of which are isoenzymes, such as SldAB1 and SldAB2 of *G. oxydans* WSH-003 (35), or are often associated with a broad range of substrates such as GoKR (11). Hence, the use of gene knockout strategy to identify the functions of some dehydrogenases of *G. oxydans*, especially the numerous unknown dehydrogenases of *G. oxydans* WSH-003, may not be appropriate. Therefore, in the present study, we expressed numerous unknown dehydrogenases of *G. oxydans* WSH-003 in *Escherichia coli* BL21 (DE3) cells and purified the products by one-step affinity chromatography with Ni-NTA agarose column to identify their functions. The results revealed a new xylitol dehydrogenase (NAD-dependent xylitol dehydrogenase 2) that could convert sorbitol to fructose. Kinetics analysis of the novel enzyme revealed some unique traits that were quite different from the known xylitol dehydrogenases. The optimum temperature and pH of the identified xylitol dehydrogenase 2 was 57°C and 12, respectively. This novel enzyme provides new insights into *G. oxydans* dehydrogenases and could have potential applications in xylitol production.

## Results

### Gene expression and purification of the identified dehydrogenase

The selected dehydrogenase from *G. oxydans* WSH-003 was successfully expressed and purified. Sequence analysis revealed that the purified enzyme, annotated as xylitol dehydrogenase 2, contained a NAD(P)-binding motif and a classical active site motif belonging to the short-chain dehydrogenase family. SDS-PAGE analysis showed an expected single band with a molecular weight of about 38 kDa (Fig. 1A), which was consistent with the calculated molecular mass based on the deduced amino acid sequence (36.6 kDa). The optimum pH and temperature for the purified xylitol dehydrogenase 2 were determined to be pH 12 (50 mM glycine-NaOH buffer) and 57°C, respectively (Fig. 1B, C), which are different from those for known xylitol dehydrogenases.

**Fig. 1.**
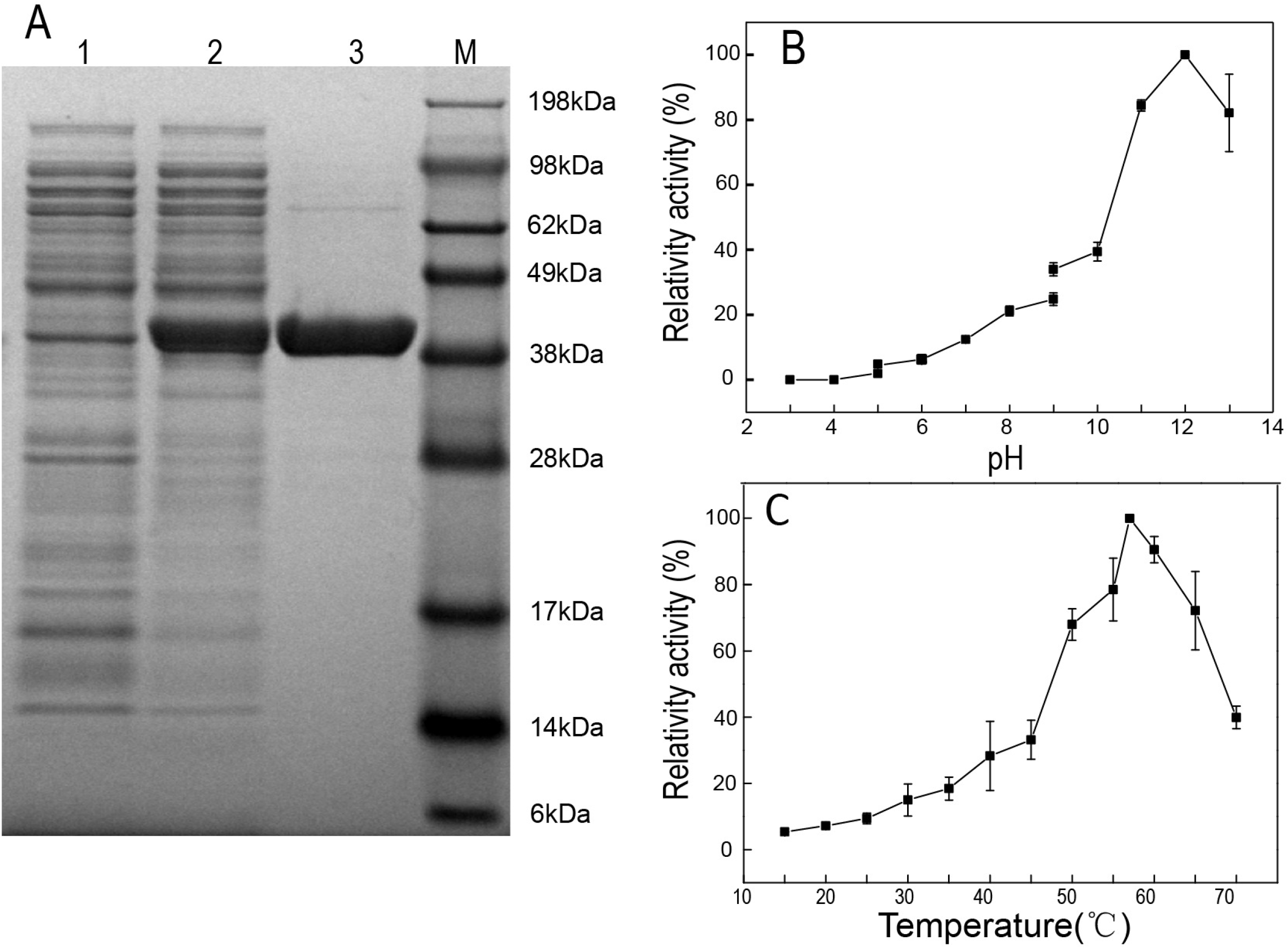
Optimum pH and temperature for xylitol dehydrogenase 2. SDS-PAGE of the identified xylitol dehydrogenase 2 purified from E. coli BL21 (DE3) cells containing pET-28a-XDH. Lane 1: E. coli BL21 containing pET-28a after induction for 16 h at 20°C. Lane 2: Recombinant strain E. coli BL21 containing pET-28a-XDH after induction for 16 h at 20°C. Lane 3: Purified recombinant enzyme. Lane M: Molecular mass markers. (B) Effect of pH on the activity of purified xylitol dehydrogenase 2. (C) Effect of temperature on the activity of purified xylitol dehydrogenase 2.

### Identification of cofactor of xylitol dehydrogenase 2

In general, dehydrogenases require some cofactors as electron acceptor, such as NAD(P), FAD/FMN, or PQQ. Most of the previously identified membrane dehydrogenases from *G. oxydans* have been reported to utilize PQQ or FAD as the cofactor. According to the prediction of transmembrane domains, xylitol dehydrogenase 2 from *G. oxydans* WSH-003 was noted to lack transmembrane domain. Therefore, the cofactor of the identified dehydrogenase was verified by using the purified enzyme to catalyze reactions with different cofactors. The results showed that xylitol dehydrogenase 2 was highly specific for NAD^+^, and no detectable enzyme activity was observed with NADP^+^, FAD, or PQQ as the cofactor (Fig. 2).

**Fig. 2.**
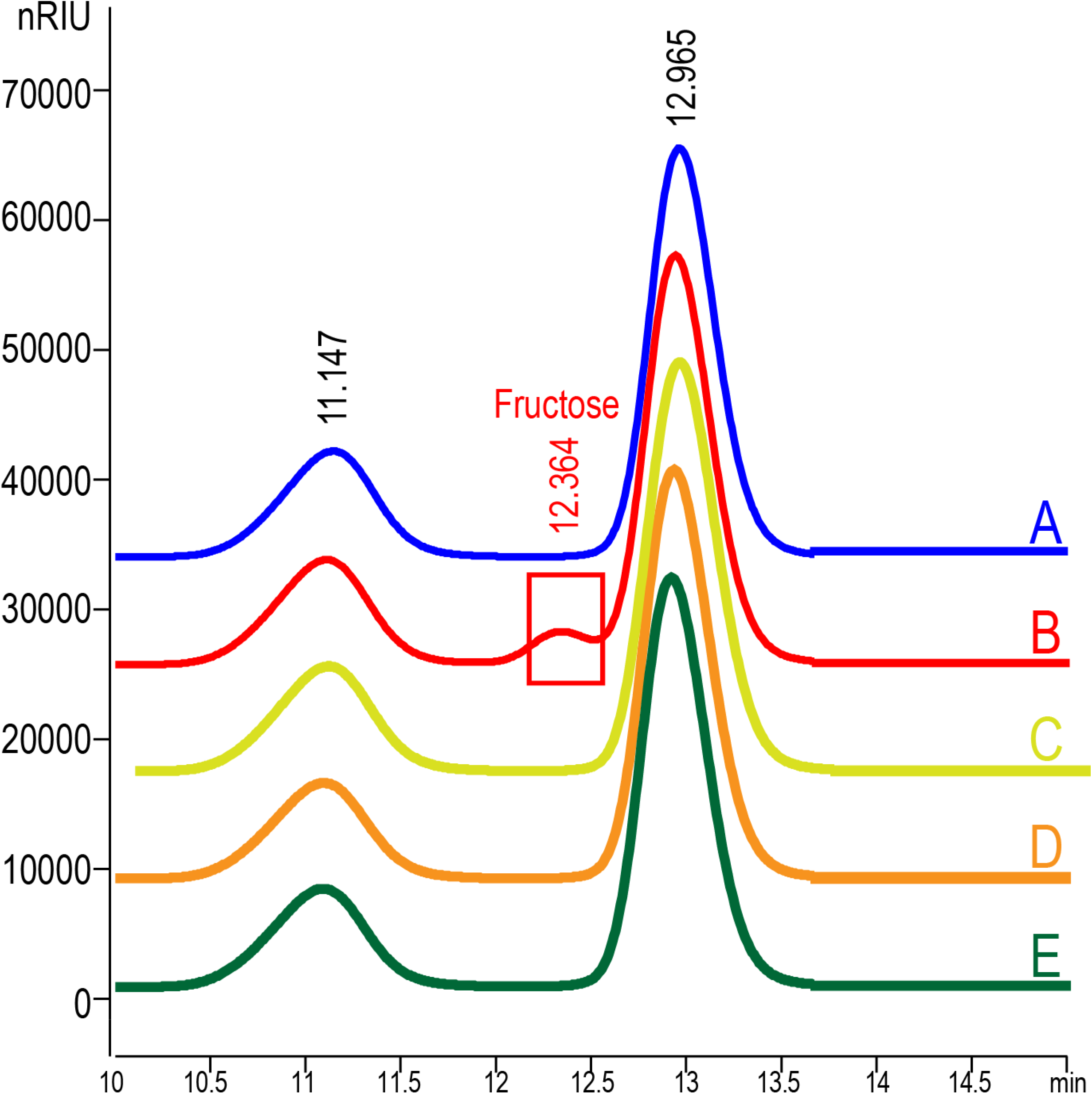
Determination of cofactor of xylitol dehydrogenase 2. Catalytic reaction of purified xylitol dehydrogenase 2 (A) without cofactor, (B) with NAD^+^, (C) with NADP^+^, (E) with FAD, and (E) with PQQ.

### Effect of EDTA and metal ions on enzyme activity

To determine the effects of chelator and metal ions on NAD-dependent xylitol dehydrogenase 2, EDTA and various ions (0.5 mM Ca^2+^, Mg^2+^, Cu^2+^, Fe^2+^, Zn^2+^, Co^2+^, Ni^2+^, Mn^2+^, Cr^3+^, and Fe^3+^) were respectively added to the reaction system. EDTA elicited no obvious effect on NAD-dependent xylitol dehydrogenase 2, indicating that the enzyme does not require chelator for its activity. However, the enzyme could be activated by Zn^2+^, Co^2+^, and Mn^2+^, among which Zn^2+^ improved the enzyme activity by 1.8 times. In contrast, Cu^2+^ could almost completely inhibit the activity of NAD-dependent xylitol dehydrogenase 2 (Fig. 3), while the rest of the examined metal ions had no obvious impact on the enzyme activity.

**Fig. 3.**
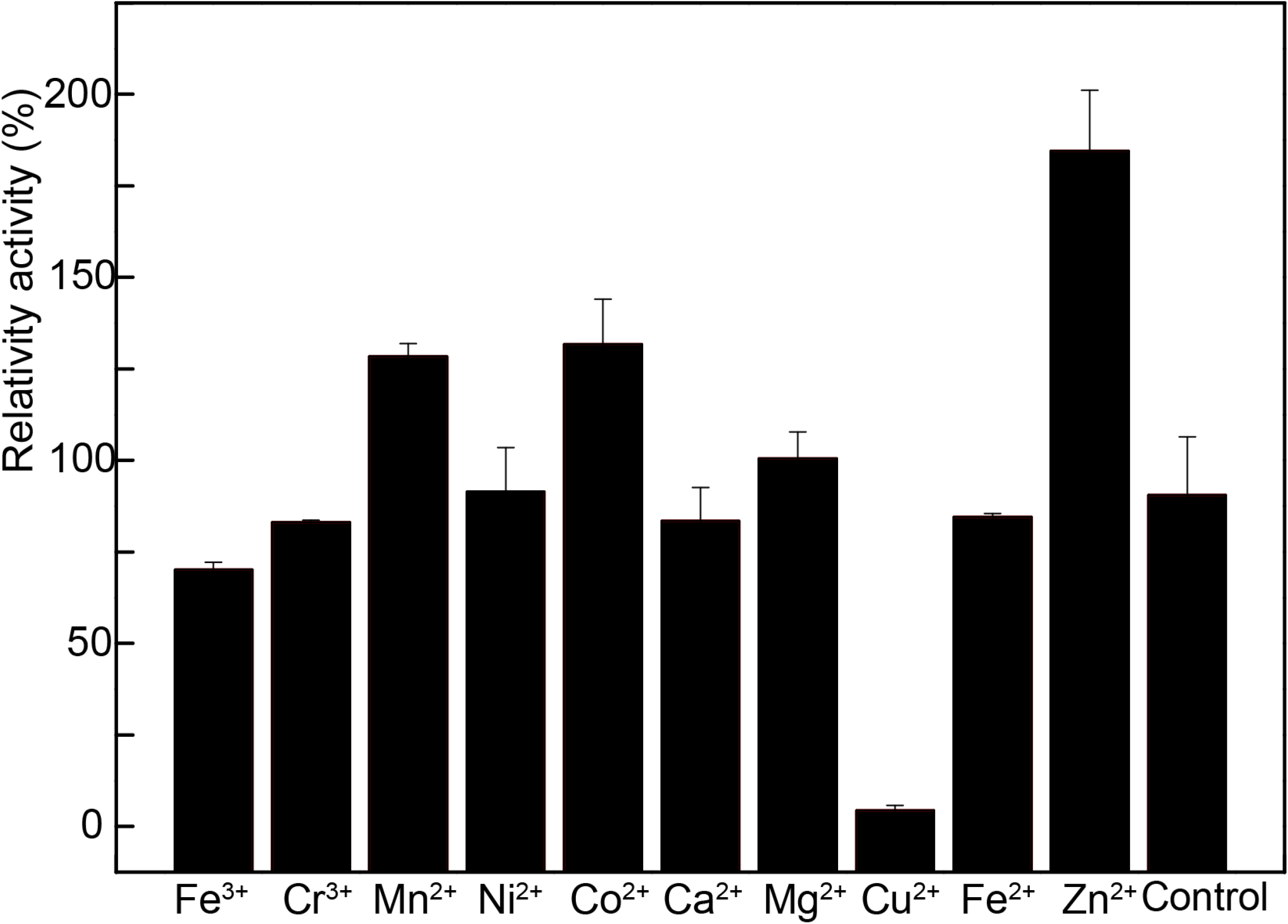
Effect of metal ions on the activity of NAD-dependent xylitol dehydrogenase 2. Relative activities of the enzyme in the presence of various metal ions, when compared with the control without metal ions.

### Substrate specificity and kinetic constants

In recent years, xylitol dehydrogenase has been used for the industrial production of xylitol, and under enhanced NADH supply, NAD-dependent xylitol dehydrogenase can reduce D-xylulose to desired xylitol. In the present study, substrate specificity analysis of NAD-dependent xylitol dehydrogenase 2 revealed that the enzyme was highly specific towards D-sorbitol and xylitol, but showed limited activity towards D-mannitol, sorbose, and glycerol. Moreover, the enzyme showed no activity when glucose, inositol, galactose, sorbitol, mannose, rhamnose, xylose, fructose, glucuronic acid, glucolactone, 2-KLG, gluconic, propanol, isopropanol, methanol, and ethanol were used as substrate (Fig. 4). To determine the kinetic constants, the initial velocities of the enzyme were determined in glycine-NaOH buffer (pH 12) with D-sorbitol (at increasing concentrations from 1 to 500 mM) under standard assay conditions, and the *K*_*m*_ and *V*_*max*_ were noted to be 4.92 μM and 196.08 μM/min, respectively.

**Fig. 4.**
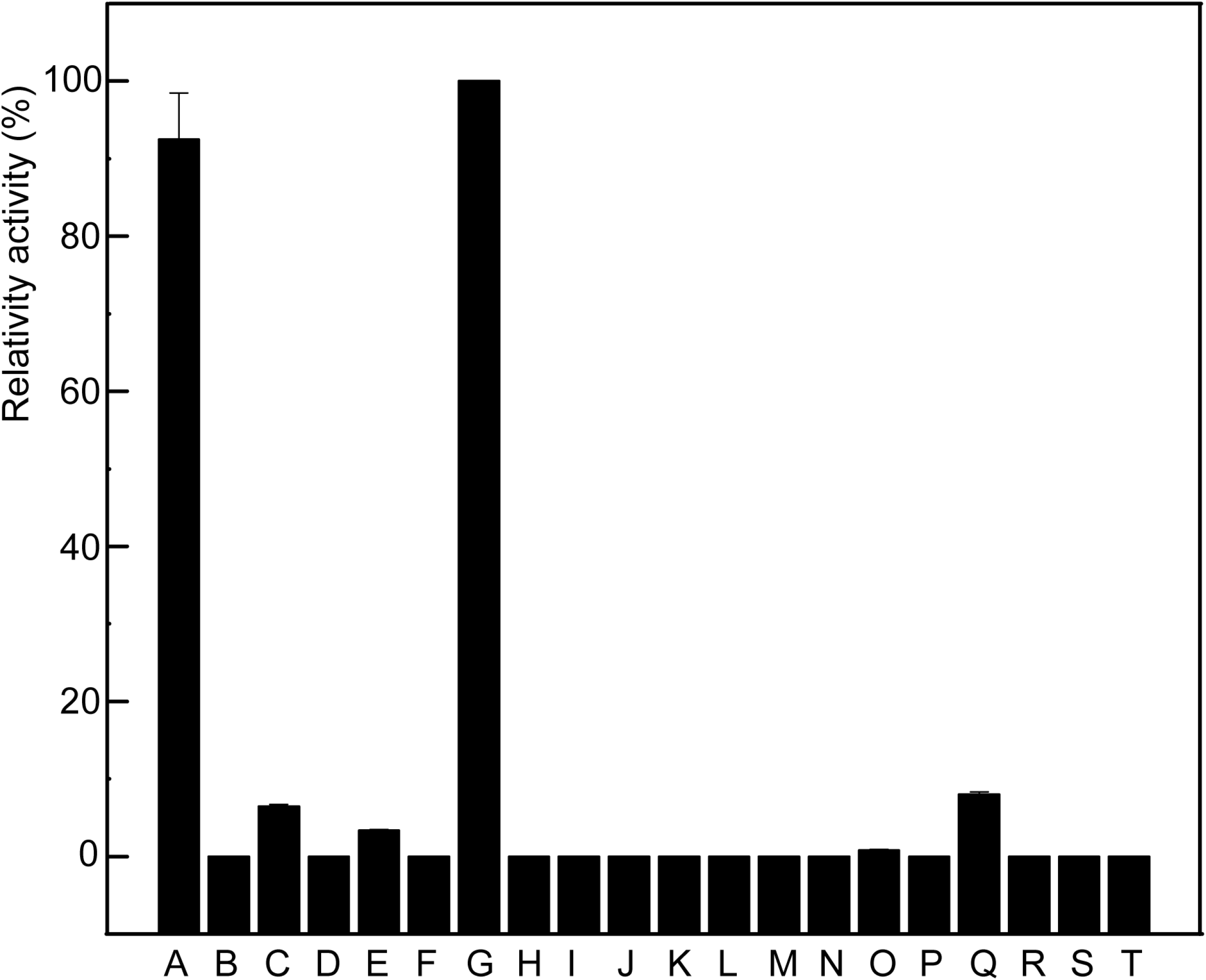
Substrate specificity of NAD-dependent xylitol dehydrogenase 2. Relative enzyme activity towards (A) xylitol, (B) glucose, (C) D-mannitol, (E) sorbose, (F) galactose, (G) sorbitol, (H) mannose, (I) rhamnose, (J) xylose, (K) fructose, (L) glucuronic acid, (M) glucolactone, (N) 2-KLG, (O) gluconic acid, (P) propanol, (Q) glycerol, (R) inopropanol, (S) methanol, and (T) ethanol.

## Discussion

In this study, a xylitol dehydrogenase from 44 uncharacterized dehydrogenases of *G. oxydans* WSH-003 was identified and characterized. This novel NAD-dependent xylitol dehydrogenase 2 could convert D-sorbitol to D-fructose, indicating certain correlation of this enzyme with pentose phosphate pathway (31). The optimum temperature and pH for the identified xylitol dehydrogenase 2 revealed its unique characteristics, when compared with some of the previously identified xylitol dehydrogenases. It has been reported that D-fructose is the major byproduct formed during the conversion of D-sorbitol to L-sorbose by *G. oxydans* in industrial-scale vitamin C production (36), and that knockout of genes involved in D-fructose production can further improve the conversion rate of D-sorbitol toL-sorbose. Owing to its unique characteristics, the NAD-dependent xylitol dehydrogenase 2 identified in the present study can be applied for the production of D-xylitol (12). To characterize all the dehydrogenases of *G. oxydans* WSH-003, the enzymes were predicted and heterologously overexpressed in *E. coli* BL21 (DE3) cells. Then, the expressed dehydrogenases were purified by one-step affinity chromatography with Ni-NTA agarose column. While most of the dehydrogenases with obvious expression levels in *E. coli* showed no activities, NAD-dependent xylitol dehydrogenase 2 could efficiently convert D-sorbitol to D-fructose.

Previous studies have indicated that majority of the numerous dehydrogenases in *G. oxydans* are membrane-bound, PQQ- or FAD-dependent enzymes with more than one subunit; for example, alcohol dehydrogenases have three subunits (37), aldehyde dehydrogenases have two subunits (38), D-sorbitol dehydrogenases have one or three subunits (18, 21, 39), and polyol dehydrogenase have two subunits (40). Most of the cytoplasmic soluble polyol dehydrogenases are NADP-dependent with more than one subunit; for instance, NADP-dependent D-sorbitol dehydrogenase have four subunits, NADP-dependent D-sorbitol dehydrogenase have two subunits (41), and NAD-dependent ribitol dehydrogenase have four subunits (42). However, the xylitol dehydrogenase 2 identified in the present study was noted to be NAD-dependent with only one subunit. The amino acid sequence of the NAD-dependent xylitol dehydrogenase 2 showed similarity to those of the enzymes in the MDR superfamily. However, the optimum pH and temperature for the oxidation activity of the NAD-dependent xylitol dehydrogenase 2 were observed to be slightly higher than those reported in earlier studies for the same reaction of xylitol dehydrogenases isolated from different strains of *G. oxydans* (43). The reason for this variation in the optimum pH and temperature for xylitol dehydrogenase activity could be owing to the different source strains from which the enzymes were isolated.

With regard to the substrate specificity of xylitol dehydrogenases, xylitol dehydrogenase from *G. oxydans* ATCC 621 has been noted to present higher catalytic activity towards sorbitol and xylitol (44), whereas xylitol dehydrogenase from *G. thailandicus* CGMCC1.3748 has been demonstrated to exhibit catalytic activity towards xylitol, D-sorbitol, D-mannitol, and D-fructose (43). Besides, while most of the known xylitol dehydrogenases have been reported to be dependent on cofactor NAD^+^, an NADP^+^-dependent xylitol dehydrogenase has been found to increase ethanol production from xylose in recombinant *Saccharomyces cerevisiae* though protein engineering (45).

In most of the identified *G. oxydans* strains, glycolysis and citric acid cycle are incomplete owing to the lack of phosphofructokinase and succinate dehydrogenase (29), which is the main reason for the low biomass yield of *G. oxydans*, when compared with other common bacteria, and a major limitation to the use of *G. oxydans* whole cell biotransformation. It has been reported that pentose phosphate pathway and ED pathway are the main catabolic routes for biomass and energy supply in *Gluconobacter* strains (46). Despite its industrial application for several decades, the metabolic pathways and regulatory mechanisms of *Gluconobacter* spp. are not yet fully elucidated (47–49). To improve the biomass of *G. oxydans*, Krajewski et al. knocked out the membrane-bound glucose dehydrogenase and soluble glucose dehydrogenase, and improved the biomass by 271% (50). An understanding of the mechanisms of catabolism of polyols, sugars, and sugar derivatives into the pentose phosphate pathway is essential for increasing the biomass and catalysis efficiency of *G. oxydans* strains. As D-sorbitol and L-sorbose cannot directly enter into the pentose phosphate pathway, they must be catabolized via some intermediates. The NAD-dependent xylitol dehydrogenase 2 identified in the present study can catalyze D-sorbitol to D-fructose, which can directly enter the pentose phosphate pathway through phosphorylation, suggesting that overexpression of this enzyme may increase the biomass of *G. oxydans* by utilizing more D-sorbitol.

In conclusion, a novel NAD-dependent xylitol dehydrogenase 2 from *G. oxydans* WSH-003 was identified in this study. Owing to its unique characteristics, such as optimum pH and temperature, the identified dehydrogenase could be used in the production of xylitol or fructose, or in regeneration of cofactor under specific conditions. Although *G. oxydans* WSH-003 has been mutated from wild-type strain at least 90 times by different methods with reliable records to improve L-sorbose production and tolerance to saccharides and alditols such as L-sorbose and D-sorbitol, generation of D-fructose as the byproduct of the strain could not be resolved. However, knockout of xylitol dehydrogenase and similar dehydrogenases could facilitate further increase in the yield of D-sorbitol to L-sorbose, which could be important for the current industrial-scale production of vitamin C.

## Materials and methods

### Genes, plasmids, and strains

The vector pMD19-T Simple and pET-28a(+) were used for vector construction and protein expression, respectively. *E. coli* JM109 cells were employed for plasmid construction and *E. coli* BL21 (DE3) cells were used for protein expression. The dehydrogenase gene (GenBank Accession No.: 29878874) was PCR-amplified from the genomic DNA of *G. oxydans* WSH-003 using the primer pair CCG<u>**GAATTC**</u>ATGGCTCAAGCTTTGGTTCTGGAAC/CCG<u>**CTCGAG**</u>TCAGCCT GGAAGCTTAATTTGTAGCTTC, purified, digested, and inserted into *Eco*RI/*Xho*I sites of pET-28a(+) to obtain pET-28a-XDH. The recombinant plasmid pET-28a-XDH was transformed into *E. coli* BL21 (DE3) cells for protein expression. All the sequences were verified by Sanger sequencing (Sangon Biotech, Shanghai, China). The transmembrane domains of the protein were predicted by using TMHMM (http://www.cbs.dtu.dk/services/TMHMM/).

### Gene expression and purification of dehydrogenase

The recombinant strain was cultured in 250-mL shake flasks containing 25 mL of Terrific broth (TB) medium. After growth to log phase (OD_600_=0.6), the cells were pre-cooled to 20°C. Then, 0.5 mM isopropyl-β-D-thiogalactopyranoside (IPTG) was added to induce protein expression, and the cells were incubated at 20°C for another 16 h for protein expression.

Subsequently, the cells were collected by centrifugation at 5,000 rpm for 5 min, washed twice with binding buffer (50 mM phosphate buffer), and lysed by sonication at 4°C. The lysate was centrifuged for 20 min at 7,000 rpm at 4°C to obtain a clear supernatant. The supernatant was passed through a 0.45-μm filter, and then applied to a 5-mL nickel-charged Hi-Trap column pre-equilibrated with binding buffer. The column was washed with 15 mL of binding buffer and then with washing buffer (50 mM phosphate buffer, 150 mM NaCl, and 50 mM imidazole; pH adjusted to 7.0) until no more protein was eluted. The column was eluted with 20 mL of eluting buffer (50 mM phosphate buffer, 150 mM NaCl, and 500 mM imidazole), and the pH of the eluent was adjusted to 7.0. The fractions were combined and dialyzed against dialysis buffer (50 mM phosphate buffer).

### Enzyme assay and identification of cofactor

The enzyme activity was measured by determining the increase in absorbance of NADH at 340 nm. The reaction mixture contained 2 mM NAD^+^, 20 mM sorbitol, 50mM phosphate buffer (pH 12), and enzyme solution to a total volume of 200 μL. One unit of enzyme activity was defined as the amount of enzyme catalyzing the formation of 1 μmol of reduced NAD^+^ per minute at 30°C under the given conditions.

### Effect of metal ions and EDTA

In order to determine the effect of the metal ions and the EDTA on the enzyme, various metal ions (0.5 mM) and EDTA (5 mM) were added individually to the reaction mixture. Relative activity was used to investigate, while the reaction mixture without any additional treatment served as a control (100%).

### Substrate specificity and determination of kinetic constants

Substrate specificity of the identified dehydrogenase was tested using 20 mM xylitol, glucose, D-mannitol, inositol, sorbose, galactose, sorbitol, mannose, rhamnose, xylose, fructose, glucuronic acid, glucolactone, 2-KLG, gluconic acid, propanol, glycerol, inopropanol, methanol, and ethanol in the above-mentioned buffers. For kinetics experiments, the substrate concentrations were varied between 1 and 500 mM and the cofactor concentration was 2 mM.

### Determination of optimum temperature and pH for the identified dehydrogenase

To determine the optimum pH, the enzyme activity was assessed in a pH range of 3–13 in the following buffers (50 mM): NaAc-HAc (pH 3.0–5.0), PBS (pH 5.0–9.0) Tris-HCl (pH 9.0–10.0), and glycine-NaOH (pH 9.0–13). Similarly, the optimal temperature for the identified dehydrogenase was determined by analyzing the enzyme activity from 20°C to 70°C.

## Acknowledgements

This work was supported by grants from the National Natural Science Foundation of China (Key Program, 31830068), the National Science Fund for Excellent Young Scholars (21822806), the Fundamental Research Funds for the Central Universities (JUSRP51701A), the National First-class Discipline Program of Light Industry Technology and Engineering (LITE2018-08), the Distinguished Professor Project of Jiangsu Province, and the 111 Project (111-2-06).

## References

1. Deppenmeier U, Hoffmeister M, Prust C. 2002. Biochemistry and biotechnological applications of *Gluconobacter* strains. Appl. Microbiol. Biotechnol. 60:233–242.

2. Giridhar R, Srivastava AK. 2000. Model based constant feed fed-batch L-sorbose production process for improvement in L-sorbose productivity. Chem. Biochem. Eng. Q. 14:133–140.

3. Yang X-P, Wei L-J, Lin J-P, Yin B, Wei D-Z. 2008. A membrane-bound PQQ-dependent dehydrogenase in *Gluconobacter oxydans* M5, responsible for production of 6-(2-hydroxyethyl) amino-6-deoxy-L-sorbose. Appl. Microbiol. Biotechnol. 74 5250–5253.

4. Poljungreed I, Boonyarattanakalin S. 2017. Dihydroxyacetone production by *Gluconobacter frateurii* in a minimum medium using fed ‐ batch fermentation. J. Appl. Chem. Biotechnol. 92:2635–2641.

5. Li K, Mao X, Liu L, Lin J, Sun M, Wei D, Yang S. 2016. Overexpression of membrane-bound gluconate-2-dehydrogenase to enhance the production of 2-keto-D-gluconic acid by *Gluconobacter oxydans*. Microb. Cell Fact. 15:121

6. Siemen A, Kosciow K, Schweiger P, Deppenmeier U. 2018. Production of 5-ketofructose from fructose or sucrose using genetically modified *Gluconobacter oxydans* strains. Appl. Microbiol. Biotechnol. 102:1699–1710.

7. Rabenhorst J, DR., Gatfield I, DR., Hilmer J-M, DR. 2001-02-28 2001. Natural, aliphatic and thiocarboxylic acids obtainable by fermentation and a microorganism therefore EP1078990 patent EP1078990.

8. Bertokova A, Bertok T, Filip J, Tkac J. 2015. *Gluconobacter* sp cells for manufacturing of effective electrochemical biosensors and biofuel cells. Chem. Pap. 69:27–41.

9. Macauley S, McNeil B, Harvey LM. 2001. The genus *Gluconobacter* and its applications in biotechnology. Crit. Rev. Biotechnol. 21:1–25.

10. Schenkmayerova A, Bertokova A, Sefcovicova J, Stefuca V, Bucko M, Vikartovska A, Gemeiner P, Tkac J, Katrlik J. 2015. Whole-cell *Gluconobacter oxydans* biosensor for 2-phenylethanol biooxidation monitoring. Anal. Chim. Acta 854:140–144.

11. Chen R, Liu X, Wang J, Lin J, Wei D. 2015. Cloning, expression, and characterization of an anti-Prelog stereospecific carbonyl reductase from *Gluconobacter oxydans* DSM2343. Enzyme Microb. Technol. 70:18–27.

12. Qi X, Zhang H, Magocha TA, An Y, Yun J, Yang M, Xue Y, Liang S, Sun W, Cao Z. 2017. Improved xylitol production by expressing a novel D-arabitol dehydrogenase from isolated *Gluconobacter sp*. JX-05 and co-biotransformation of whole cells. Bioresour. Technol. 235:50–58.

13. Zhu J, Xie J, Wei L, Lin J, Zhao L, Wei D. 2018. Identification of the enzymes responsible for 3-hydroxypropionic acid formation and their use in improving 3-hydroxypropionic acid production in *Gluconobacter oxydans* DSM 2003. Bioresour. Technol. 265:328–333.

14. Matsushita K, Toyama H, Adachi O. 1994. Respiratory chains and bioenergetics of Acetic Acid Bacteria. Adv. Microb. Physiol. 36:247–301.

15. Matsushita K, Yakushi T, Toyama H, Adachi O, Miyoshi H, Tagami E, Sakamoto K. 1999. The quinohemoprotein alcohol dehydrogenase of *Gluconobacter suboxydans* has ubiquinol oxidation activity at a site different from the ubiquinone reduction site. Biochim. Biophys. Acta, Bioenerg. 1409:154–164.

16. Schweiger P, Volland S, Deppenmeier U. 2007. Overproduction and characterization of two distinct aldehyde-oxidizing enzymes from *Gluconobacter oxydans* 621H. J. Mol. Microbiol. Biotechnol. 13:147–155.

17. Shinagawa E, Matsushita K, Toyama H, Adachi O. 1999. Production of 5-keto-D-gluconate by acetic acid bacteria is catalyzed by pyrroloquinoline quinone (PQQ)-dependent membrane-bound D-gluconate dehydrogenase J. Mol. Catal. B: Enzym. 6:341–350.

18. Choi ES, Lee EH, Rhee SK. 1995. Purification of a membrane-bound sorbitol dehydrogenase from *Gluconobacter suboxydans*. FEMS Microbiol. Lett. 125:45–49.

19. Kim T-S, Patel SKS, Selvaraj C, Jung W-S, Pan C-H, Kang YC, Lee J-K. 2016. A highly efficient sorbitol dehydrogenase from *Gluconobacter oxydans* G624 and improvement of its stability through immobilization. Sci. Rep. 6: 33438.

20. Shibata T, Ichikawa C, Matsuura M, Takata Y, Noguchi Y, Saito Y, Yamashita M. 2000. Cloning of a gene for D-sorbitol dehydrogenase from *Gluconobacter oxydans* G624 and expression of the gene in *Pseudomonas putida* IFO3738. J. Biosci. Bioeng. 89:463–468.

21. Sugisawa T, Hoshino T. 2002. Purification and properties of membrane-bound D-sorbitol dehydrogenase from *Gluconobacter suboxydans* IFO 3255. Biosci., Biotechnol., Biochem. 66:57–64.

22. Asakura A, Hoshino T. 1999. Isolation and characterization of a new quinoprotein dehydrogenase, L-sorbose/L-sorbosone dehydrogenase. Biosci., Biotechnol., Biochem. 63:46–53.

23. Saito Y, Ishii Y, Hayashi H, Imao Y, Akashi T, Yoshikawa K, Noguchi Y, Soeda S, Yoshida M, Niwa M, Hosoda J, Shimomura K. 1997. Cloning of genes coding for L-sorbose and L-sorbosone dehydrogenases from *Gluconobacter oxydans* and microbial production of 2-keto-L-gulonate, a precursor of L-ascorbic acid, in a recombinant *G. oxydans* strain. Appl. Environ. Microbiol. 63:454–460.

24. Zahid N, Deppenmeier U. 2016. Role of mannitol dehydrogenases in osmoprotection of *Gluconobacter oxydans*. Appl. Microbiol. Biotechnol. 100:9967–9978.

25. Adachi O, Toyama H, Matsushita K. 1999. Crystalline NADP-dependent D-mannitol dehydrogenase from *Gluconobacter suboxydans*. Bioscience Biotechnology and Biochemistry 63:402–407.

26. Vangnai AS, Promden W, De-Eknamkul W, Matsushita K, Toyama H. 2010. Molecular characterization and heterologous expression of quinate dehydrogenase gene from *Gluconobacter oxydans* IFO3244. Biochemistry (Moscow) 75:452–459.

27. Yakushi T, Komatsu K, Matsutani M, Kataoka N, Vangnai AS, Toyama H, Adachi O, Matsushita K. 2018. Improved heterologous expression of the membrane-bound quinoprotein quinate dehydrogenase from *Gluconobacter oxydans*. Protein Expression Purif. 145:100–107.

28. Yakushi T, Terada Y, Ozaki S, Kataoka N, Akakabe Y, Adachi O, Matsutani M, Matsushita K. 2018. Aldopentoses as new substrates for the membrane-bound, pyrroloquinoline quinone-dependent glycerol (polyol) dehydrogenase of *Gluconobacter* sp. Appl. Microbiol. Biotechnol. 102:3159–3171.

29. Prust C, Hoffmeister M, Liesegang H, Wiezer A, Fricke WF, Ehrenreich A, Gottschalk G, Deppenmeier U. 2005. Complete genome sequence of the acetic acid bacterium *Gluconobacter oxydans*. Nat. Biotechnol. 23:195–200.

30. Greenfield S, Claus GW. 1972. Nonfunctional tricarboxylic acid cycle and the mechanism of glutamate biosynthesis in *Acetobacter suboxydans*. J. Bacteriol. 112:1295–1301.

31. Kiefler I, Bringer S, Bott M. 2017. Metabolic engineering of *Gluconobacter oxydans* 621H for increased biomass yield. Appl. Microbiol. Biotechnol. 101:5453–5467.

32. Peters B, Junker A, Brauer K, Mühlthaler B, Kostner D, Mientus M, Liebl W, Ehrenreich A. 2013. Deletion of pyruvate decarboxylase by a new method for efficient markerless gene deletions in *Gluconobacter oxydans*. Appl. Microbiol. Biotechnol. 97:2521–2530.

33. Peters B, Mientus M, Kostner D, Junker A, Liebl W, Ehrenreich A. 2013. Characterization of membrane-bound dehydrogenases from *Gluconobacter oxydans* 621H via whole-cell activity assays using multideletion strains. Appl. Microbiol. Biotechnol. 97:6397–6412.

34. Mientus M, Kostner D, Peters B, Liebl W, Ehrenreich A. 2017. Characterization of membrane-bound dehydrogenases of *Gluconobacter oxydans* 621H using a new system for their functional expression. Appl. Microbiol. Biotechnol. 101:3189–3200.

35. Gao L, Zhou J, Liu J, Du G, Chen J. 2012. Draft genome sequence of *Gluconobacter oxydans* WSH-003, a strain that is extremely tolerant of saccharides and alditols. J. Bacteriol. 194:4455–4456.

36. Macauley-Patrick S, McNeil B, Harvey LM. 2005. By-product formation in the D-sorbitol to L-sorbose biotransformation by *Gluconobacter suboxydans* ATCC 621 in batch and continuous cultures. Process Biochem. 40:2113–2122.

37. Adachi O, Tayama K, Shinagawa E, Matsushita K, Ameyama M. 1978. Purification and characterization of particulate alcohol dehydrogenase from *Gluconobacter suboxydans*. Agric. Biol. Chem. 42:2045–2056.

38. Adachi O, Tayama K, Shinagawa E, Matsushita K, Ameyama M. 1980. Purification and characterization of membrane-bound aldehyde dehydrogenase from *Gluconobacter suboxydans*. Agricultural and Biological Chemistry 44:503–515.

39. Shinagawa E, Matsushita K, Adachi O, Ameyama M. 1982. Purification and characterization of D-sorbitol dehydrogenase from membrane of *Gluconobacter suboxydans* var. α. Agric. Biol. Chem. 46:135–141.

40. Matsushita K, Fujii Y, Ano Y, Toyama H, Shinjoh M, Tomiyama N, Miyazaki T, Sugisawa T, Hoshino T, Adachi O. 2003. 5-keto-D-gluconate production is catalyzed by a quinoprotein glycerol dehydrogenase, major polyol dehydrogenase, in *Gluconobacter* species. Appl. Environ. Microbiol. 69:1959–1966.

41. Adachi O, Ano Y, Moonmangmee D, Shinagawa E, Toyama H, Theeragool G, Lotong N, Matsushita K. 1999. Crystallization and properties of NADPH-dependent L-sorbose reductase from *Gluconobacter melanogenus* IFO 3294. Biosci., Biotechnol., Biochem. 63:2137–2143.

42. Adachi O, Fujii Y, Ano Y, Moonmangmee D, Toyama H, Shinagawa E, Theeragool G, Lotong N, Matsushita K. 2001. Membrane-bound sugar alcohol dehydrogenase in acetic acid bacteria catalyzes L-ribulose formation and NAD-dependent ribitol dehydrogenase is independent of the oxidative fermentation. Biosci., Biotechnol., Biochem. 65:115–125.

43. Zhang H, Yun J, Zabed H, Yang M, Zhang G, Qi Y, Guo Q, Qi X. 2018. Production of xylitol by expressing xylitol dehydrogenase and alcohol dehydrogenase from *Gluconobacter thailandicus* and co-biotransformation of whole cells. Bioresour. Technol. 257:223–228.

44. Sugiyama M, Suzuki S, Tonouchi N, Yokozeki K. 2003. Cloning of the xylitol dehydrogenase gene from *Gluconobacter oxydans* and improved production of xylitol from D-arabitol. Biosci., Biotechnol., Biochem. 67:584–591.

45. Matsushika A, Watanabe S, Kodaki T, Makino K, Inoue H, Murakami K, Takimura O, Sawayama S. 2008. Expression of protein engineered NADP^+^-dependent xylitol dehydrogenase increases ethanol production from xylose in recombinant *Saccharomyces cerevisiae*. Appl. Microbiol. Biotechnol. 81:243–255.

46. e Muynck C, Pereira CSS, Naessens M, Parmentier S, Soetaert W, Vandamme EJ. 2007. The genus *Gluconobacter oxydans*: Comprehensive overview of biochemistry and biotechnological applications. Crit. Rev. Biotechnol. 27:147–171.

47. Klasen R, Bringer-Meyer S, Sahm H. 1992. Incapability of *Gluconobacter oxydans* to produce tartaric acid. Biotechnol. Bioeng. 40:183–186.

48. Rauch B, Pahlke J, Schweiger P, Deppenmeier U. 2010. Characterization of enzymes involved in the central metabolism of *Gluconobacter oxydans*. Appl. Microbiol. Biotechnol. 88:711–718.

49. Gupta A, Qazi GN, Verma V. 1997. Transposon induced mutation in *Gluconobacter oxydans* with special reference to its direct-glucose oxidation metabolism. FEMS Microbiol. Lett. 147:181–188.

50. Krajewski V, Simić P, Mouncey NJ, Bringer S, Sahm H, Bott M. 2010. Metabolic engineering of *Gluconobacter oxydans* for improved growth rate and growth yield on glucose by elimination of gluconate formation. Appl. Environ. Microbiol. 76:4369–4376.

